# Pheniqs: Fast and flexible quality-aware sequence demultiplexing

**DOI:** 10.1101/128512

**Authors:** Lior Galanti, Dennis Shasha, Kristin C. Gunsalus

## Abstract

1

**Motivation:** Output from high throughput sequencing instruments often exceeds what is necessary to assay a single sample. To better utilize this capacity, multiple samples are independently tagged with a unique “barcode” sequence and are then pooled, or “multiplexed”, and sequenced together. Classifying, or “demultiplexing”, the reads involves decoding the barcode sequence. Although instruments estimate the probability of incorrectly calling each nucleobase, available demultiplexers do not consult those estimates or report classification error probabilities.

**Results:** We present Pheniqs, a fast and flexible sequence demultiplexer and quality analyzer. In addition to providing an efficient implementation of the widespread *minimum distance decoder*, Pheniqs introduces a novel *Phred-adjusted maximum likelihood decoder* that consults base calling quality scores and estimates the probability of a barcode decoding error. Setting an upper bound on the permissible error provides an intuitive way to control demultiplexing confidence and directly influence precision and recall. Pheniqs supports FASTQ and multiple Sequence Alignment/Map formats and uses auxiliary SAM tags to report both library classification and demultiplexing error probability. Evaluation on both real and semi-synthetic data indicates that Pheniqs is faster than existing demultiplexers, substantially when demultiplexing longer reads, and achieves greater accuracy by correctly reflecting quality measurements.

**Availability and Implementation:** Implemented in multithreaded C++ and available under the terms of the AGPL-3.0 license agreement at http://github.com/biosails/pheniqs. Manual and examples are available at http://biosails.github.io/pheniqs.

## 2 Introduction

Sequencing platforms cycle through nucleic acid polymers one nucleobase at a time, convert technology-specific raw data formats into DNA sequence “reads”, and then report the identity of the observed nucleobase at each position along with an estimated probability of an incorrect assignment. This process, commonly referred to as “base calling”, is often bundled by commercial analysis pipelines with demultiplexing in a single black-box procedure. From both a scientific and a software engineering perspective, it is preferable to separate these two tasks since base calling is platform-dependent, but demultiplexing is not and would benefit from the transparency provided by peer review and open access to algorithms and code. Although errors introduced during library preparation or sequencing can affect base calling fidelity and hence proper barcode identification, currently available tools do not consult base calling quality scores or estimate decoding error probabilities; thus, researchers typically rely on simple heuristics to estimate library cross-contamination during downstream analysis (Yang et al., 2015). Control of classification errors in multiplexed sequence data is important both to help maximize yield, minimize sequencing costs and improve the quality of reported results, particularly for applications that require comparisons between samples such as differential gene expression analysis. Sequencing centers are usually interested in assessing instrument performance and yield, while researchers are also concerned with read assignment confidence. Quality control traditionally involves assessing Phred score distributions across cycles and along the reads, and then removing or masking potentially erroneous base calls and technical sequences in a subsequent step.

Ideally, a general purpose tool would address the basic requirements for functionality and be designed around a consistent API to simplify integration with other utilities; however, traditional tools that are the current standard in high-throughput sequence analysis fall short of this ideal in various ways. The fastx toolkit has not been updated since early 2010 and produces only very basic analysis in comma separated format; fastqc produces more elaborate analysis but is mostly human readable and requires more work to integrate the analysis into a pipeline; and Rqc will only operate on a subset of the reads (10% by default) to mitigate R’s sluggish IO handling. Recently developed tools like MultiQC (Ewels et al., 2016) create an integrated report and can visualize results from multiple tools and across many samples, making global trends and biases more easily apparent. Such tools are designed to be extended and can benefit from an efficient analysis engine that collects statistics during demultiplexing and makes them available in an easily consumable format.

With regard to base calling and demultiplexing, Illumina’s bcl2fastq tool follows different workflows for different instruments and relies on poorly documented and untunable quality assessment procedures. The Picard toolkit from the Broad Institute provides a much more uniform approach for base calling that applies to all Illumina sequencers, but similarly relies on a *minimum distance decoder*. Both bcl2fastq and Picard can be challenging to integrate into a pipeline and are exclusively geared toward demultiplexing Illumina data. Other standalone demultiplexers include the C program fastq-multx (Aronesty, 2013), fastx barcode splitter (a perl script and part of the FASTX-toolkit) and the Java package Je (Girardot et al., 2016). However fastq-multx does not allow for addressing fragments of the read and so requires preprocessing to demultiplex when the barcode is not exactly present in a separate file. fastx barcode splitter takes a single FASTQ (Cock et al., 2010) from standard input and so can only demultiplex single-end reads with an inline barcode on either the 5’ or 3’ end and is significantly slower than other tools. Je is a wrapper around several Picard tools that enables direct demultiplexing of FASTQ files. It offers additional flexibility for locating barcodes and the ability to extract and place directly flanking molecular barcodes (UMI or Unique Molecular Identifier) into the FASTQ read identifier comment. None of those tools consults the base calling quality scores, provides confidence estimates or supports SAM metadata manipulation. Since no generic standardized tool that can easily be integrated into pipelines is available to handle unconventional experimental designs, sequencing centers often deliver multiplexed FASTQ files, forcing researchers to rely on their own home-brewed demultiplexers that are often inefficient, overly simplistic and potentially buggy.

Pheniqs **(PH**ilology **EN**coder w**I**th **Q**uality **S**tatistics, pronounced “phoenix”) overcomes the limitations of other available tools described above. Pheniqs can read, write and manipulate FASTQ files as well as the *Sequence Alignment/Map format* (Li, Handsaker, et al., 2009) encoded in a SAM file or one of its binary compressed variants BAM and CRAM. Using a powerful yet simple syntax, Pheniqs can extract fragments of read segments by addressing either the 5’ end, 3’ end or both. The fragments are then used to construct output template segments, multiplex barcodes and/or molecular barcodes. This generic approach will scale well as experimental designs continue to evolve. Since Pheniqs analyzes read quality during demultiplexing, when the reads are already present in computer memory, quality assessment incurs only a negligible overhead. Moreover, Pheniqs produces a quality report of both inputs and outputs that exhaustively consults all the reads and is encoded in JSON, which can be easily consumed by many programming languages. We propose that making available an efficient, Open Source and peer-reviewed implementation that can handle unconventional experimental designs and the latest file formats will greatly benefit the field (Eklund, Nichols, and Knutsson, 2016). Pheniqs thus provides an efficient, Open Source implementation that can handle unconventional experimental designs and the latest file formats; in addition it enables demultiplexing error probabilities to percolate down the analysis pipeline, making them available for future hypothesis testing.

## 3 Approach

Pheniqs offers two demultiplexing strategies: the traditional *minimum distance decoder*, previously described in detail in the context of sequencing (Mir et al., 2013), and a novel *Phred-adjusted maximum likelihood decoder* that not only outperforms the former but also consults base calling quality scores and reports a decoding error probability. The *minimum distance decoder* bundled with Pheniqs can produce results identical to traditional demultiplexers and relies on the minimal pair-wise mismatch distance between the barcode sequences (defined as the number of corresponding positions with non-identical bases) to provide a worst-case scenario estimate for an unambiguous error correction threshold (Shannon, 1948). Pheniqs will automatically establish the maximum number of mismatches that each defined barcode set can tolerate when using *minimum distance decoding* by inspecting the pairwise barcode Hamming distance matrix. It will also report the matrix, which can be insightful when designing a barcode set, during input validation.

The *Phred-adjusted maximum likelihood decoder* uses an error model that takes into account the Phred-encoded base calling error probabilities provided by the sequencing instrument. By examining the match probability of the read segment spanning an expected barcode position with each multiplexing barcode in a given set, it takes advantage of the uneven barcode mismatch distance distribution to improve recovery, while flagging as ambiguous any demultiplexing decisions supported primarily by low quality cycles. Match probability estimates provide a classification confidence for each read
— much the same way that mapping quality, introduced by BWA (Li and Durbin, 2009), provides an estimate for a mapping confidence. Setting a threshold on the confidence provides a reportable upper bound for the error probability of cross-library read contamination for each sample, an essential measurement in some experimental designs.

## 4 Methods

### 4.1 Theoretical Model

Demultiplexing involves extracting an observed subsequence *r* from the read and decoding *b*, the barcode sequence, from *r*. Let *r* ∈ {*A, C, G, T, N*}^*n*^ be an observed sequence of length *n* extracted from the read and 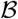 a given set of non-identical barcodes from {*A, C, G, T, N*}^*n*^ of size 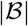 identifying the multiplexed samples. A decoder is denoted as a decision function 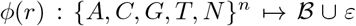 where ε denotes a decoding failure. The probability that *r* is observed given that *b* was sequenced is *P*(*r*|*b*) and the probability that *b* was sequenced given *r* was observed is *P*(*b*|*r*),

Intuitively, we want a maximum likelihood estimator 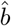, so a good decoder should select a 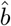 which maximizes the posterior probability

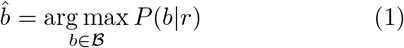

Applying Bayes’ rule to **Equation 1** gives

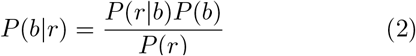

*P*(*b*) is the expected fraction of the pooled sample identified by *b* after accounting for the rate of occurrence of foreign sequences *P*_ε_, and *P*(*r*) is independent of decoding and given by the *law of total probability*

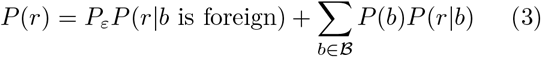

Since foreign sequences are assumed to produce random observations *P*(*r*|*b* is foreign) = 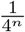 and

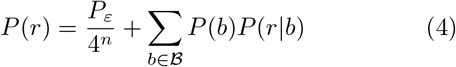

**Equation 2** can be written as

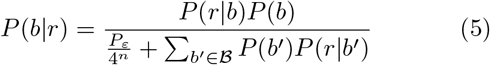
Since the denominator of **Equation 2** is constant for any given *r* and 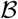, for the purpose of estimating 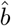 it is sufficient to solve

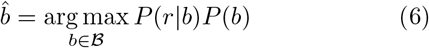
which is also faster and more numerically stable.

### 4.2 Phred-adjusted maximum likelihood decoding

Our novel *Phred-adjusted maximum likelihood decoder* solves **Equation 6** by directly estimating *P*(*r*|*b*) for each 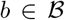 from the Phred scores and additionally uses the sample pooling composition and foreign sequence probability as estimators for the priors *P*(*b*). Once 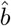 has been identified it proceeds to compute 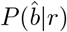 using **Equation 5** so that the decision function ϕ(*r*) becomes

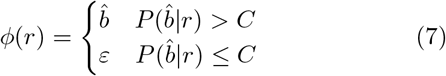
where *C* is a user provided confidence threshold for the minimum acceptable probability of a correct decoding. To compute *P*(*r*|*b*) from the Phred score, assuming errors on base calls are unrelated, we take the product over the bases (Edgar and Flyvbjerg, 2015)

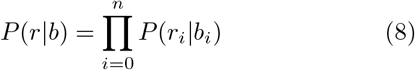
where *p*(*r*_*i*_|*b*_*i*_) is defined to be

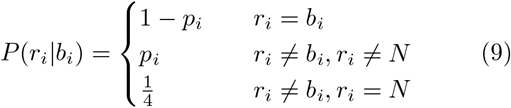
and *p*_i_ is the base calling error probability for base call *r*_*i*_ decoded from the Phred value *q*_*i*_ by applying 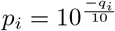 for each position *i* in *r*.

Estimating 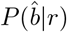 directly enjoys the benefit that the confidence of classifying *r* as 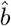 is a highly desirable reportable statistic and the probability of a decoding error becomes

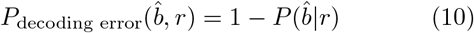

Phred is a log-scaled encoding and as such loses accuracy as the values becomes smaller. When all the probabilities produced by **Equation 8** are extremely small the 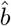 produced by **Equation 6** can be misleading. This can occur when the observed sequence *r* is relatively high quality but very poorly matches all barcodes in 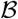, or when the overall quality of the barcode is very low. To control for such cases, we introduce a second threshold that compares 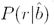 to the probability of observing a random sequence of length n:

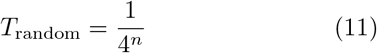

Reads with 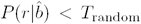 should probably be considered a decoding failure without further consideration since the initial evidence supporting their classification is inferior to that provided by a random sequence.

Directly estimating 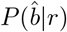 allows Pheniqs to report a demultiplexing quality for every read. Exposing a tunable threshold *C* allows researchers to choose between assignment confidence and yield depending on the application, and to factor the reported confidence into downstream analysis. This high precision estimation of the assignment probability takes advantage of the non-uniform pairwise Hamming distance distribution between the pooled barcodes to recover additional reads that would otherwise be considered a decoding failure by *minimum distance decoders*. Factoring the instrument-reported error probabilities provides better control for false classifications when critically informative bases on the barcode have low confidence.

### 4.3 Minimum distance decoding

By contrast, we show that the assumptions made by *minimum distance decoding* lead to inaccurate estimates of *P*(*b*|*r*) when applied to sequencing. If samples are pooled uniformly, than 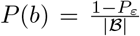 for all 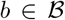 and **Equation 5** is simplified to

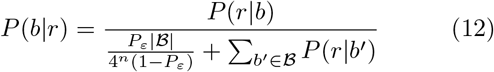

Further minimum distance decoding assumes that *P*_*ε*_ → 0 and 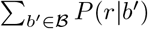 is constant, so *P*(*b*|*r*) is proportional to *P*(*r*|*b*) and the same 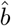 that solves **Equation 1** also solves

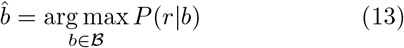
which is often referred to in coding theory as the *maximum likelihood decoding rule*. Minimum distance decoding solves **Equation 13** by minimizing the Hamming distance to the observed sequence r:

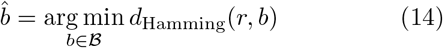
where the Hamming distance *d*_Hamming_(*r, b*) between *r* and *b* is defined as the number of corresponding positions with non-identical bases. A lower bound on the number of correctable substitution errors is given by the seminal 1948 mathematical theory of communication (Shannon, 1948) to be

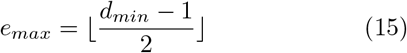
where *d*_*min*_ is the minimum pairwise barcode Hamming distance and 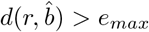 indicates a decoding failure, so the decision function ϕ(*r*) becomes

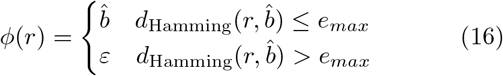
Currently available demultiplexers use *minimum distance decoding* and rely on **Equation 14** coupled with a user provided threshold, even though any value higher than e_max_ is not guaranteed to yield unambiguous assignments in any non-trivial scenario. The *minimum distance decoder* implemented within Pheniqs takes the extra step to compute and use e_max_ as an upper bound to the distance threshold for *minimum distance decoding*.

*Minimum distance decoding* is suboptimal, as it ignores many properties specific to sequencing. First, it relies on **Equation 7** to decode 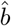, so assumes a uniform pooling concentration and ignores *P*_ε_, the presence of foreign sequences in the solution (such as *PhiX174* virus, which is used by all illumina sequencers for instrument calibration and can sometimes constitute as much as 50% of total DNA in applications with low expected base diversity). Second, it ignores instrument-reported error probabilities and computes a biased estimator for *P*(*r*|*b*), effectively assuming the confidence is always 1. Third, it uses *e*_*max*_ but that is an overly pessimistic threshold for declaring a decoding failure: when the pairwise Hamming distance distribution is uneven, as often occurs, it results in an uneven increase in the number of decoding failures across the barcode set in a way that can bias studies that rely on quantification.

## 5 Implementation

Pheniqs can demultiplex sequencing reads with barcodes located at arbitrary locations. Pheniqs extracts fragments of read segments by addressing either the 5’ end, 3’ end or both and optionally reverse complements them. It can then use the fragments to construct both output template segments, multiplex barcodes, and/or molecular barcodes, providing a completely generic approach that can accommodate any potential barcoding scheme and experimental design. Pheniqs performs an exhaustive quality assessment during demultiplexing and emits a report suitable for further integration. Pheniqs can read, write, and manipulate FASTQ files as well as the commonly used *Sequence Alignment/Map* (SAM) file format or one of its binary compressed variants BAM and CRAM. It can annotate SAM encoded files with user-specified Read Group metadata corresponding to the multiplexed libraries. The facilities provided by HTSlib allow demultiplexing results to be written to a single, smaller file and at the same time support richer annotation, which significantly simplifies pipelines. Pheniqs reports the decoded barcode and its corresponding quality scores in the standardized *BC* and *QT* SAM auxiliary tags. It also reports the estimated error statistics in two new SAM auxiliary fields, which we propose to be standardized in the SAM specification: *DQ*, the demultiplexing error probability, and *EE*, the expected number of erroneous bases (Edgar and Flyvbjerg, 2015). Inclusion of these statistics in a standard report format will enable users to easily assess the distribution of demultiplexing quality scores across sample datasets. The ability to both consume from and produce to a single file, coupled with flexible read layout manipulation, enables Pheniqs to be used with POSIX standard streams.

**Figure 1:**
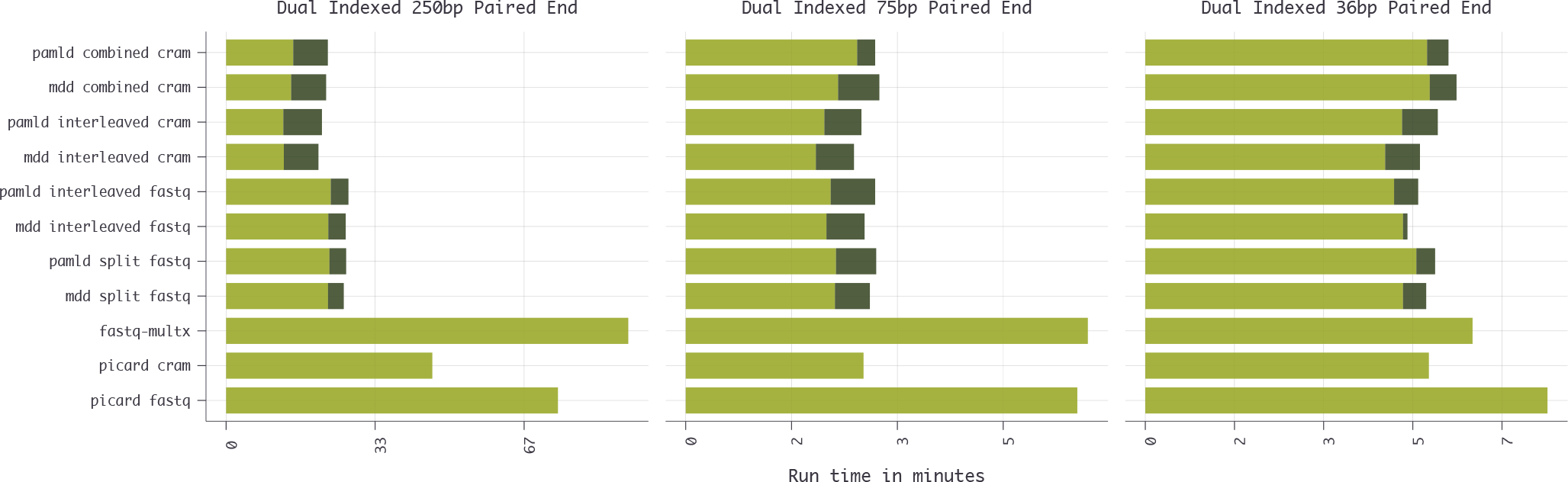
Runtime benchmarks (supplementary Table 1) comparing Picard IlluminaBasecallsToFastq, Picard IlluminaBasecallsToSam, fastq-multx, and several Pheniqs configurations. Dark stacked bars represent the additional processing time when collecting quality statistics during demultiplexing. Three lanes were used: 1. a dual-indexed paired-end HiSeq 2500 RapidRun v2 flowcell with a 250,8,8,250 configuration containing 182,198,212 reads and 21 multiplexed libraries; 2. a dual-indexed paired-end NextSeq 500 flowcell with a 75,8,8,75 configuration containing 42,681,647 reads and 48 multiplexed libraries; 3. a dual-indexed paired-end NextSeq 500 flowcell with a 36,9,9,36 configuration containing 93,209,096 reads and 96 multiplexed libraries. Benchmarking was conducted on a dual socket Intel Xeon E5-2650 v2 @ 2.60GHz for a total of 16 hyper-threaded cores and 64GB of RAM running Ubuntu 14.04.5 with linux kernel 3.13.0-101. Kernel virtual memory was flushed before every run.

This can significantly reduce disk usage or IO limitation in some applications while at the same time allow researchers to more interactively experiment with different confidence thresholds.

## 6 Discussion and results

### 6.1 Benchmarking and comparing to other tools

Pheniqs can arrange read segments in arbitrary layouts. To be able to compare its performance to other tools we benchmarked three commonly used layouts: **split, interleaved,** and **combined** (Figure 1 and supplementary Table 1). We refer to a layout as **split** when each segment of the read in each multiplexed library is written to a separate file; as **interleaved** when all segments are written consecutively to the same file but different multiplexed libraries are still written to separate files; and as **combined** when all segments of all reads in all libraries are written to the same file.

For a dual indexed paired end configuration Pheniqs requires all 4 raw read segments with their corresponding quality scores in either FASTQ or HTSlib formats. The Picard *IlluminaBasecallsToFastq* command was used to generate 4 multiplexed FASTQ files as input for pheniqs. All lanes were base called using Picard version 2.7.1.

Results of demultiplexing with the Picard *IlluminaBasecallsToFastq* and *IlluminaBasecallsToSam* commands are reported here for completeness but are difficult to compare since both commands take raw Illumina intensity files and perform base calling and demultiplexing in a single step. The Picard demultiplexer otherwise relies on a standard *minimum distance decoder* that does not consider the Phred error probabilities except for optionally masking base calls below a user-specified threshold. *IlluminaBasecallsToFastq* could not produce **interleaved** FASTQ files, *IlluminaBasecallsToSam* could not produce a **combined** output, and *fastq-multx* only produces **split** FASTQ files. *fastx barcode splitter* and *Je* were not included in the benchmark but are both reported (Girardot et al., 2016) to be 17 and 4.5 times respectively slower than *fastq-multx*. Pheniqs is slightly faster than both *fastq-multx* and *Picard* when processing short reads but is 2.7 times faster when processing 250 nucleotide paird-end reads in FASTQ format and 3 times faster when processing CRAM.

### 6.2 Comparing PAMLD and MDD with biological data

Comparing the accuracy of different demultiplexers is challenging for biological data since the ground truth is unknown. Here we show how to create an approximation to the ground truth and use that to compare methods.

We used Pheniqs to demultiplex 366,573,792 paired-end reads with 2 36-nucleotide segments and 2 8-nucleotide index segments from an Illumina NextSeq 500 flowcell with both decoders (Figure 2). The reads contained wild isolates of the *Saccharomyces cerevisiae, Naumovia castelli*, and *Candida orthopsilosis* yeast strains. The libraries were multiplexed with two barcode sequences containing 8 nucleotides each. The *PAMLD* confidence threshold defaults to 0.99, corresponding to an error probability lower than 1%.

30 libraries containing very few reads or predominantly reads that failed to align to any of the yeast strains were removed. 286,326,757 reads from the remaining 66 libraries were broken into three sets: 281,339,717 **consensus** (98.26%), 4,980,484 **P_only_** (1.74%), and 6,556 **M**_**only**_ (0.00229%). The **consensus** set contains reads that both *MDD* and *PAMLD* successfully decoded as the same barcode. The **P_only_** set contains reads that *MDD* failed to decode but were successfully decoded by *PAMLD*. The **M_only_** set contains reads that *MDD* successfully decoded but *PAMLD* rejected and classified as undetermined. Since the majority of currently available data has been demultiplexed with *MDD*, we used the **consensus** set to estimate the ground truth. We would hope that reads from the **P_only_** set present a distribution of biological alignments similar to reads from the same library in the **consensus** set, which should be the case if they do indeed form a subset of that library, while we expect **M_only_** reads to present a dissimilar distribution.

**Figure 2:**
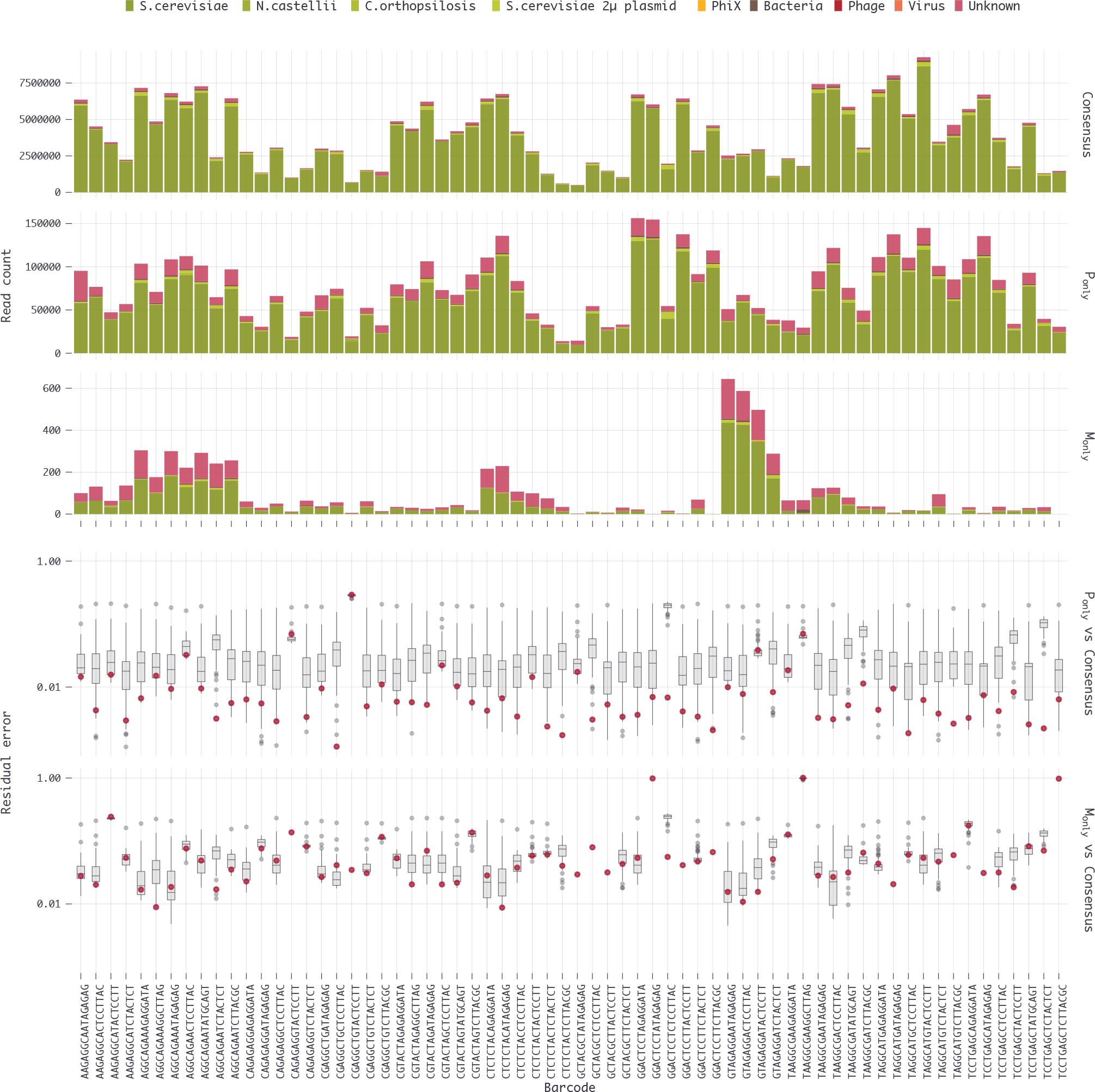
Alignment classification of 286,326,757 reads in 66 libraries. Top three rows: demultiplexed libraries classified by alignment to reference sequences and separated into three sets: 281,339,717 **consensus** (98.26%), 4,980,484 **P**_**only**_ (1.74%), and 6,556 **M**_**only**_ (0.00229%). **Consensus** and **P**_**only**_ reads are more similar in composition. Bottom two rows: the residual sum of squares, expressed as the R value, between the alignment distributions of **P**_**only**_ and **M**_**only**_ reads against **consensus** reads. Comparisons within each library (red dots) or against all other libraries (grey distributions) show that **P**_**only**_ reads generally show a stronger correlation (lower R) with **consensus** reads from the same library.

Reads were first processed with dustmasker 1.0.0 (Morgulis et al., 2006) to mask out low complexity sequences and classified as *PhiX Control, Bacteria, Virus*, and *Phage* using kraken 0.10.6 (Wood and Salzberg, 2014). The remaining reads were aligned using BWA 0.7.8-r455 to the relevant yeast references as well as the *Saccharomyces cerevisiae* 2μ plasmid. To validate the **P**_only_ and **M**_only_ sets we estimated the correlation of the their biological alignment distribution to the **consensus** sets. For each multiplexed library we computed the residual sum of squares for the distributions within the same library and between each library and all other libraries.

*Phred-adjusted maximum likelihood decoding* failures are less sensitive to base-calling errors, which easily throw off the *minimum distance decoder*. Since we expect the **P**_**only**_ and **M**_**only**_ sets to contain more reads with a higher expected error rate that are more likely to fail to align, we excluded from this computation reads that failed all alignment attempts.

All sets recovered by *PAMLD* are more highly correlated with the **consensus** set from their respective library than from one of the other libraries, while the **M**_**only**_ sets with sufficient data rarely show a better correlation with their *MDD* classification (Figure 2).

### 6.3 Comparing PAMLD and MDD with semisynthetic data

Error patterns on barcode sequences vary depending on the sequencing instrument used and its underlying error rate, the number of nucleotides in the barcode, the barcode sequence position along the read, and the library preparation protocol. Since *PAMLD* does not consider the error probabilities on different nucleotides to be related and indel errors occur at a much lower rate than substitutions, we simulated position-independent errors using a simple model based on the substitution rates observed for the same Illumina NextSeq flowcell as in Figure 2. Given that an expected nucleobase *b* ∈ {*A, C, G, T*} is sequenced with a Phred quality score *q*, we computed a maximum likelihood estimator 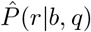 for the probability that a nucleobase *r* ∈ {*A, C, G, T, N*} is base-called by counting the relative number of times such an event was observed within the 16-nucleotide barcodes of the classified **consensus** reads.

To simulate a barcode we selected one at random from a given collection, and to simulate noise we picked a random sequence from a pool of 100,000 reads simulated from the *PhiX174* genome using the short read simulator wgsim 0.3.2. We then paired the simulated barcode sequence with the sequence of Phred quality scores for a random observed barcode from the run. 1% of the simulated reads were picked from the *PhiX174* pool while the remainder were uniformly picked from the set of known barcodes in the collection. To simulate sequencing substitution errors we iterated over the base-quality score pairs (*b, q*) and, using the distribution implied by 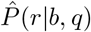, assigned a corresponding r to each position in the barcode. We simulated two datasets of 100 million reads each: one with a collection of 96 16-base barcodes and the other with a collection of 10 6-base barcodes. Each library was evaluated as a binary classifier, so a correct assignment was counted as a true positive (*TP*) while an incorrect assignment was counted as a false negative (*FN*) for the correct library and a false positive (*FP*) for the incorrectly assigned library. We then summed up the values from all libraries and computed a *false discovery rate* and *miss rate* (where *false discovery rate* is 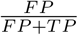 and *miss rate* is 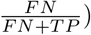). Results for *MDD* and *PAMLD* with different values for the confidence parameter are reported in Figure 3 and supplementary Table 2.

**Figure 3:**
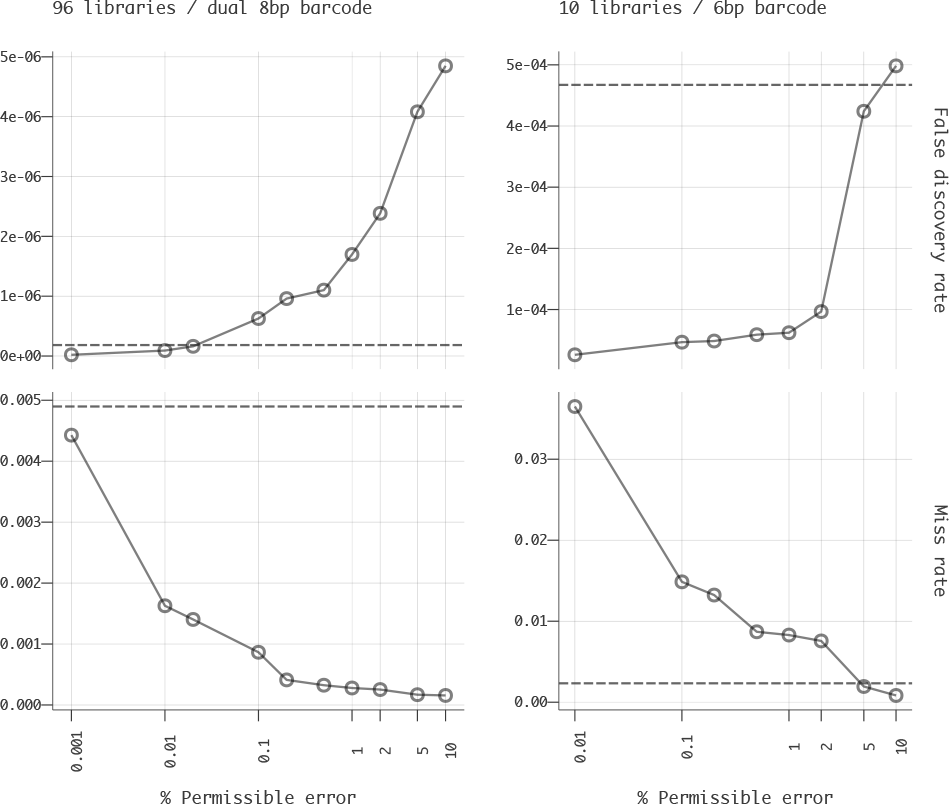
Comparison of decoding performance for *PAMLD* at varying confidence thresholds versus *MDD* (dashed line) in two demultiplexing scenarios using 100 million simulated reads. The *PAMLD* confidence parameter provides an easy way to choose between false positives and false negatives.

Six-base barcodes provide little room for error and thus false positive rates are much higher overall than for 16-base barcodes. At a confidence setting of 95%, *PAMLD* achieved a 10% improvement in *false discovery rate* while still improving *miss rate* by 20% over *MDD*. At the default 99% confidence level, *PAMLD* was able to reduce the number of false positives 7.5-fold, but at the expense of a 3.7-fold increase in false negatives. Using 16-base barcodes, false positives are sufficiently low to be considered insignificant, but any confidence threshold lower than 99.99% yielded a significant improvement in miss rate relative to *MDD*. Using the default 99% confidence yielded a 17.6-fold improvement in miss rate while adding only 150 false positive reads.

Ultimately, the choice of controlling for false positives or false negatives depends on the application, and *PAMLD* effectively provides researchers with a disciplined method to choose between the two.

### 6.4 Future work

Several additional features are planned for future software releases. These include quality-based and adapter trimming (in order to provide an integrated solution for preprocessing and quality control) as well as support for decoding of reads produced by sequencing platforms other than Illumina. Pheniqs already provides facilities to extract molecular barcodes and their corresponding quality scores into the non-standard *RX* and *QX* tags already adopted by the community, which we also recommend to be standardized; however, estimating the decoding error probability for random tags whose a priori distribution is unknown will require further development of the decoding model. Future work will expand *Phred-adjusted maximum likelihood decoding* to estimate a corrected molecular barcode and encode it in the *BX* tag together with the corresponding decoding error probability in the *PX* tag. Internally Pheniqs can already handle IUPAC ambiguity encoding (Nomencl, 1970) and support for this, by adjusting **Equation 9,** is pending further investigation. It should be noted here that the CRAM binary format, which is vastly superior to BAM both in terms of decoding and encoding speed and disk utilization, does not support IUPAC ambiguity encoding.

The comprehensive JSON-encoded quality report provides a unique opportunity to compile a composite report of reads instead of their constituent segments, as most existing tools do, which would be more intuitive for paired-end reads. Compiling the report during demultiplexing will also enable facile comparison of technical or biological replicates in different samples. A graphical tool that leverages the report produced by Pheniqs is currently in the works.

## 7 Conclusion

Pheniqs is a multithreaded, portable, efficient, robust, and flexible demultiplexer for high-throughput sequencing applications. It is highly configurable, easily integrated into existing pipelines, and offers numerous convenient features including common preprocessing tasks and versatile handling of barcode designs and input/output formats. In addition to the standard *minimum distance decoder* — the sole demultiplexing regime implemented by all other existing tools — Pheniqs uniquely offers an innovative probabilistic decoder that provides enhanced performance and introduces, for the first time, the ability to report estimates of demultiplexing error probabilities in standard output formats based on read quality scores emitted by all major sequencing platforms. Pheniqs fills a major gap in existing bioinformatics pipelines, and we anticipate that its adoption will greatly facilitate a wide range of sequence analysis applications.

## Acknowledgments

We are grateful to Alan Twaddle from the New York University Center for Genomics and Systems Biology for his support in benchmark analysis and to Giuseppe Saldi, Jillian Rowe, and Nizar Drou from the NYU Abu Dhabi Center for Genomics and Systems Biology for their insightful comments on the initial draft. We also thank Dr. David Gresham for allowing us to use data from his sequencing runs as a test case for highly multiplexed data and Or Biran and Kapil Thadani for their comments on the statistical evaluation and notation.

